# Accurate prediction of transmembrane *β*-barrel proteins from sequences

**DOI:** 10.1101/006577

**Authors:** Sikander Hayat, Chris Sander, Arne Elofsson, Debora S. Marks

## Abstract

Transmembrane *β*-barrels are known to play major roles in substrate transport and protein biogenesis in gram-negative bacteria, chloroplasts and mitochondria. However, the exact number of transmembrane *β*-barrel families is unknown and experimental structure determination is challenging. In theory, if one knows the number of strands in the *β*-barrel, then the 3D structure of the barrel could be trivial, but current topology predictions do not predict accurate structures and are unable to give information beyond the *β*-strands in the barrel. Recent work has shown successful prediction of globular and alpha-helical membrane proteins from sequence alignments, by using high ranked evolutionary couplings between residues as distance constraints to fold extended polypeptides. However, these methods, have not addressed the calculation of precise *β*-sheet hydrogen bonding that defines transmembrane *β*-barrels, and would be required to fold these proteins successfully. Hence we developed a method (EVFold_BB) that can successfully model transmembrane *β*-barrels by combining evolutionary couplings together with topology predictions. EVFold_BB is validated by the accurate all-atom 3D modeling of 18 proteins, representing all known membrane *β*-barrel families that have sufficient sequences available. To demonstrate the potential of our approach we predict the unknown 3D structure of the LptD protein, the plausibility of its accuracy is supported by the blindly predicted benchmarks, and is consistent with experimental observations. Our approach can naturally be extended to all unknown *β*-barrel proteins with sufficient sequence information.

**Significance:** EVFold_BB predicts fast, accurate 3D models of large membrane *β*-barrels that are notoriously hard to solve experimentally. The major advance is the use of evolutionary couplings from sequence alignments together with the *β*-strand prediction to ascertain accurate hydrogen bond between the*β*-strands that gives rise to the canonical barrel shapes. The method will enable biological research into outer-membrane proteins.

## Introduction

Transmembrane *β*-barrels (TMBs) constitute between 1-4% of all genes in eukaryotes and bacteria genomes [1]. Though these estimates vary substantially, due to the difficulty of the classification by sequence alone, there has been increasing interest in these proteins as their roles has been uncovered in a wide range of biomedical fields. These roles include outer-membrane-protein biogenesis [2–4], antibiotic resistance [5], vaccine design [6, 7], translocation of virulence factors [8–10], and the design of cancer therapeutics [11]. In many of these examples, the 3D structure of the TMB has been crucial in elucidating the mechanisms of, for instance, substrate diffusion [12] and voltage-gating [13] or in aiding therapeutic design [14].

Existing computational approaches can successfully identify the location of *β*-strands [15, 16], but 3D modeling techniques such as tobmodel and 3d-spot [17, 18] cannot account for the non-ideal, non-circular shape of the barrel pore nor the barrel-plug interactions. Recent work has shown that 3D structures of globular [19–21] and alpha-helical membrane proteins [22, 23] can be successfully predicted from the identification of co-evolved residues in multiple sequence alignments (MSA). The idea is that some spatially close residues co-evolve to maintain structural and functional integrity of the protein [24]. Though this approach was first reported over 20 years ago [25], only the recent approaches that use a global statistical model successfully identify sufficiently accurate close contacts from evolutionary co-variation, to fold proteins *de novo* [21, 24, 26-28]. The key innovation was to de-convolute direct from indirect correlations using maximum-entropy or related formalisms under the constraints of the data [21, 24, 26-28]. However, it is not known whether these methods can be used for large TMBs where the number of *β*-strands and their registration is critical to the overall structure. Here, we develop a hybrid method based on evolutionary couplings (EVFold-PLM [24, 27]) together with an improved *β*-strand prediction method (boctopus [15]) to generate pairs of residue restraints that can be used to fold large TMBs. The method generates accurate 3D structures of TMBs, identifies barrel and plug domain interactions and detects residues that are likely to be functionally important [23]. The *de novo* prediction of LptD, a TMB with an unknown structure and no structural homologues [29], is supported by the accuracy of the benchmark set of 18 proteins that are generated blindly to the known structures.

## Results

### Accurate folding of large membrane *β*barrels when there are sufficient sequences

We generated sequence alignments of 24 TMBs with a known structure from the OPM_PDB database [30] and selected 18 families that had enough sequences (Methods). EVFold-PLM [24, 27] was then used to calculate evolutionary coupling scores (ECs), for all pairs of residues (Fig. 1 and Methods). The proportion of accurate contacts in the top L/2 raw ECs using EVFold-PLM is ∼ 48% (Table 1), where L is the length of the protein. For residues on adjacent transmembrane *β*-strands, the average number of correctly predicted contacts predicted by EVFold-PLM is ∼62% (Table 1).

**Fig. 1.**
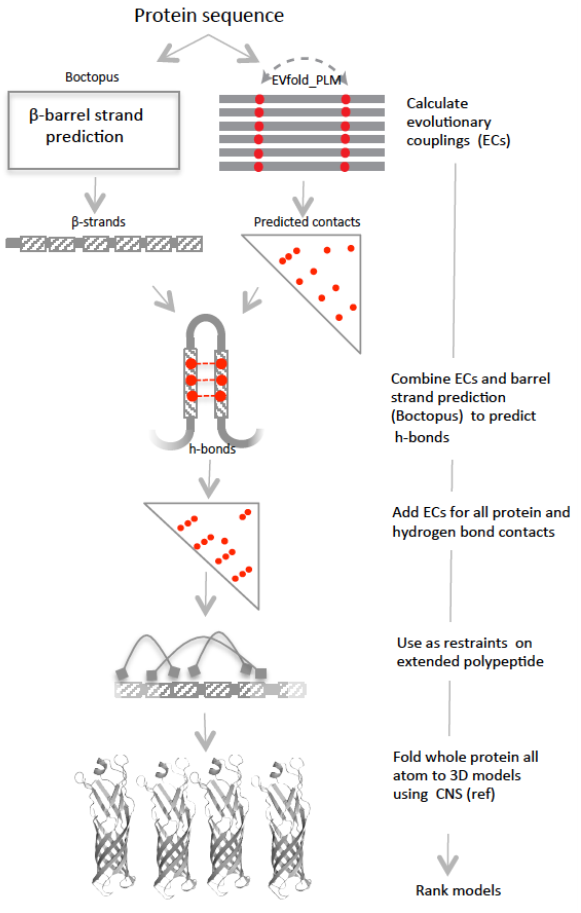
EVFold_BB pipeline to *de novo* fold TMBs. EVFold-PLM is used to generate ECs from a MSA of a protein. β-strand location is predicted using boctopus2.0. ECs and β-strand location is used to determine the strand-registration by shifting adjacent strands up/down +/− 3 residues with respect to each other. Configuration that satisfies most ECs is chosen. Hydrogen bonds are placed on residue pairs that are in register such that dyad repeat pattern and right-hand twist is maintained. Inferred hydrogen bond constraints and other non strand-strand are used to *de novo* fold TMBs. Generated models are blindly ranked.

**Table 1–.**
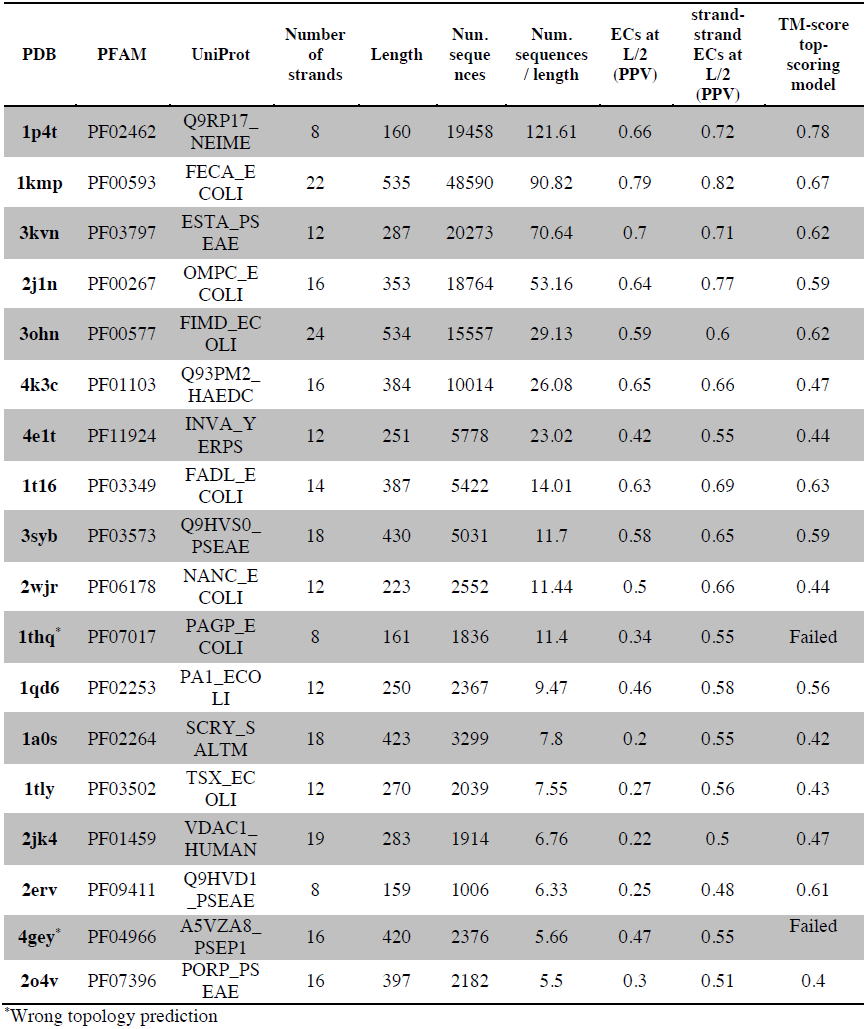
Benchmark results

**Table 1–.** Extended benchmark results

| PDB | Oligo<br>meric<br>state | Meff | Filter SS<br>clashes<br>ECs at<br>L/2<br>(PPV) | Filter SS<br>clashes -<br>strand-strand ECs<br>at L/2 (PPV) | Hbonds<br>in top H<br>(ECs<br>alone) | Hbonds<br>in top H<br>(strand<br>shift +<br>ECs) | Hbonds<br>(strand<br>shift +<br>ECs)<br>PPV | TM-score<br>top-scoring<br>model | TM-score<br>best<br>possible<br>model |
| --- | --- | --- | --- | --- | --- | --- | --- | --- | --- |
| **1p4t** | M | 4402.1 | 0.75 | 0.72 | 14 | 33 | 0.69 | 0.78 | 0.83 |
| **1kmp** | M | 19162.26 | 0.8 | 0.82 | 30 | 82 | 0.63 | 0.67 | 0.7 |
| **3kvn** | M | 2952.05 | 0.75 | 0.71 | 16 | 42 | 0.58 | 0.62 | 0.68 |
| **2j1n** | T | 3432.5 | 0.68 | 0.77 | 21 | 57 | 0.58 | 0.59 | 0.67 |
| **3ohn** | D | 1678.46 | 0.63 | 0.6 | 29 | 65 | 0.44 | 0.62 | 0.62 |
| **4k3c** | M | 3704.72 | 0.74 | 0.66 | 30 | 46 | 0.48 | 0.47 | 0.52 |
| **4e1t** | M/D? | 388.58 | 0.46 | 0.55 | 14 | 21 | 0.29 | 0.44 | 0.47 |
| **1t16** | M | 1916.85 | 0.71 | 0.69 | 11 | 54 | 0.64 | 0.63 | 0.73 |
| **3syb** | M | 1049.11 | 0.61 | 0.65 | 12 | 46 | 0.42 | 0.59 | 0.69 |
| **2wjrr** | M | 186.85 | 0.59 | 0.66 | 18 | 35 | 0.5 | 0.44 | 0.51 |
| **1thq\*** | M | 120.03 | 0.38 | 0.55 | NA | NA | NA | Failed | Failed |
| **1qd6** | M/D | 357.43 | 0.47 | 0.59 | 14 | 32 | 0.44 | 0.56 | 0.61 |
| **1a0s** | T | 370.93 | 0.28 | 0.56 | 19 | 39 | 0.35 | 0.42 | 0.43 |
| **1tly** | M | 268.27 | 0.4 | 0.53 | 9 | 29 | 0.4 | 0.43 | 0.46 |
| **2jk4** | M/D | 948.73 | 0.29 | 0.5 | 11 | 40 | 0.33 | 0.47 | 0.54 |
| **2erv** | M | 436.41 | 0.29 | 0.48 | 4 | 24 | 0.49 | 0.61 | 0.65 |
| **4gey\*** | M | 850.54 | 0.55 | 0.55 | NA | NA | NA | Failed | Failed |
| **2o4v** | T | 1219.51 | 0.37 | 0.51 | 13 | 16 | 0.16 | 0.4 | 0.43 |

\*Wrong topology prediction, M – monomeric, D – dimer, T – trimeric, Meff – Effective number of sequences in alignment

For 16 of the 18 proteins in our benchmark dataset, the correct number and approximate location of *β*-strands can be predicted using boctopus (see Methods and [15]). In addition, contact maps based on ECs from EVFold-PLM overlap with the predicted*β*-strand locations (Fig. 2 and Fig. S1). However, folding *β*-barrels with these sparse, and sometimes incorrect constraints produces inaccurate barrel geometry (data not shown). Although 9 of the 18 benchmark proteins have no high ranking ECs between the first and last strands (Fig. S1), the missing contacts in 7 of these proteins are plausibly due to lack of sequence coverage in the alignment. Fig. S2). Since we know that the membrane*β*-strands follow a strict hydrogen bond repeat pattern across adjacent strands, it makes sense to estimate the complete hydrogen-bond pattern (Methods), in order to fold TMBs more accurately. Briefly, the algorithm to determine the *β*-strands hydrogen bonds, starts with the top L adjacent strand-strand ECs, and the predicted*β*-strand positions (Methods), shifts the pairwise strands to find the optimal registration for each strand pair such that the ECs score between the two strands is maximized. The algorithm identifies 661/897 (∼ 73%) hydrogen bond pairs observed in 237 adjacent *β*-strand pairs across 16 proteins for which correct topology was predicted by boctopus2.0 (Fig. 2).

**Fig. 2.**
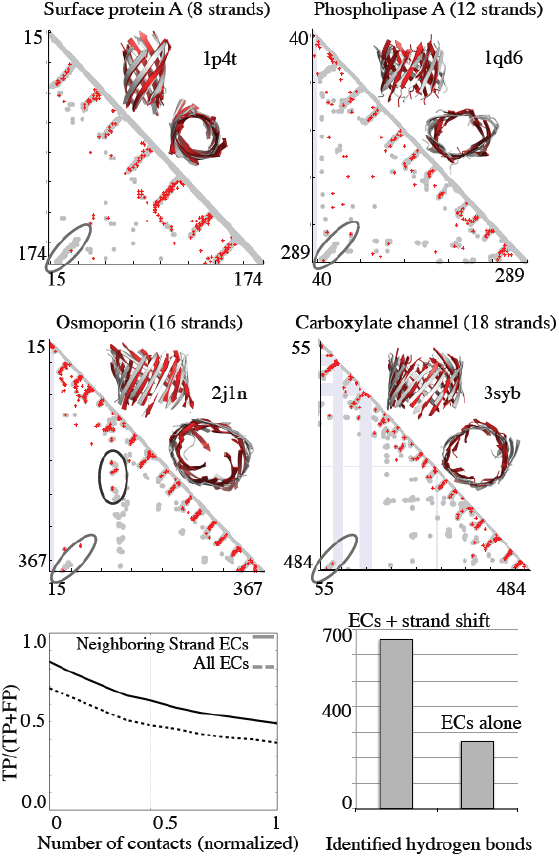
*de novo* predicted 3D models of transmembrane β-barrels with known structure. Contact maps (red - ECs, gray - crystal contacts <= 5 Å, blue - gaps in crystal structure) and front and top view of folded structures (red - *de novo* folded, gray - crystal structure) for 4 proteins in dataset. Bottom left panel - fraction of correct ECs over total predictions (solid), adjacent strand-strand (dashed) normalized to L - protein length. Right panel - Number of correct hydrogen bonds predicted using boctopus+ECs and ECs alone at length H, H - number of hydrogen bonds predicted by boctopus+ECs.

Hydrogen bond distances are then combined with the top ranked non-adjacent strand-strand ECs as well as loop ECs, obtained directly from EVFold-PLM to generate distance constraints on extended polypeptides that are then folded in CNS [31] as described previously [24]. The template modelling (TM-score) of top ranked models ranges from 0.40 to 0.78 (Table 1 and Supplementary Data). Although 12 proteins have a TM score of > 0.5 for the best-generated model (Methods), models with TM score of > 0.5 can only be identified for 9 proteins by our ranking procedure (Table 1). For the other 7, the top-ranked models have TM score in the range 0.4 to 0.47, most probably because these proteins have fewer sequences in the alignments (TSX_ECOLI, PORP_PSEAE, VDAC1_HUMAN and SCRY_SALTM have < 10 sequences / residue), have a low number of diverse sequences in the alignment (INVA_YERPS, NANC_ECOLI, SCRY_SALTM and TSX_ECOLI have less than 500 sequences in the alignment after redundancy reduction), or have multimer signals that have not been removed (PORP_PSEAE, VDAC1_HUMAN and SCRY_SALTM) (Table 1).

In addition to folding TMBs, we identify interactions between *β*-barrels and plug domains. For instance, the FecA barrel domain consists of 22 transmembrane *β*-strands along with a large plug domain (∼126 residues) and 9 out of 10 high-ranking ECs identify accurate contacts between these two distinct domains (Fig. 3). The TM-score for the top-ranking FecA model with barrel alone and barrel+plug domain is 0.67 and 0.68, respectively (Supplementary Data). This shows that EVFold_BB can identify co-evolved residues between two interacting domains and generate a 3D model for the entire protein.

**Fig. 3.**
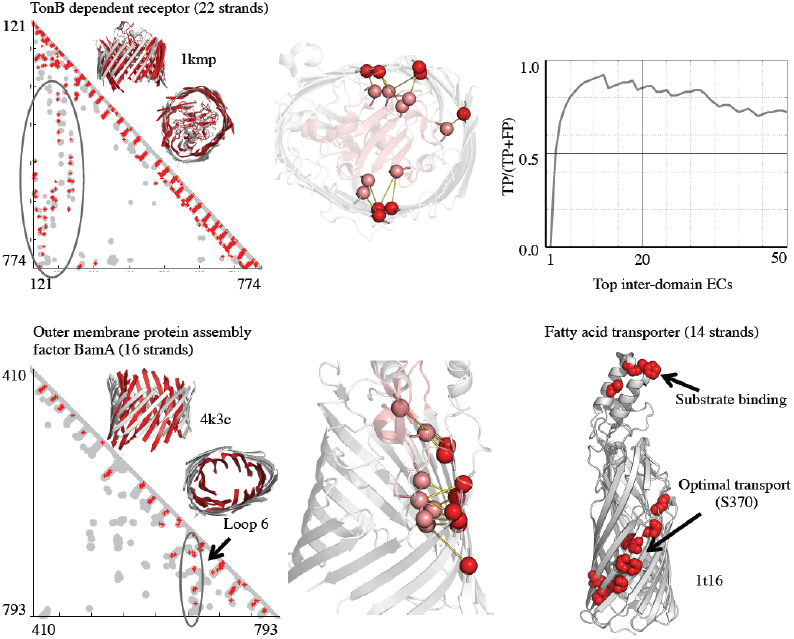
ECs predict barrel/non-barrel interactions and reveal functional contacts. FecA - Interactions between the barrel (245–774) and plug (121–244) domain in FecA are highlighted in the predicted contact map (red - ECs, gray - crystal contacts <= 5 Å). Top 10 inter-domain contacts between the barrel (red) and plug domain (pink) are shown on crystal structure and have a PPV of 0.9. BamA - Top ranking interactions between the extracellular loop 6 and the barrel domain are circled in the contact map and are shown on the crystal structure (ECs on barrel - red, on Loop 6- pink). FadL - Top 10 residues with most EC interactions (in top L/2) are shown in red on crystal structure (gray). These residues include the binding site in loops 3 and 4 on the extracellular side and S370, which is required for optimal long-chain fatty acid transport.

### A 3D model of the unknown structure of LptD

We generated 3D models of the outer membrane protein LptD and propose residues that might play a role in lipopolysaccharide (LPS) transport [29]. The LptD sequence consists of a N-terminus cytosolic domain and a C-terminus *β*-barrel domain. Based on experimental evidence, the C-terminal barrel domain has been suggested to start from residue 200 [29]. The LptD barrel domain has no detectable sequence similarity to proteins with any known structure. Three different prediction methods (pred-tmbb [32], proftmb [33] and boctopus2.0 (Methods)) are used to predict *β*-strands in the putative barrel domain (residue 200-784). Boctopus predicts 26 strands and the other two methods predict 24 strands. These methods provide consensus on the location of 20 strands but do not agree on the predicted location of the other *β*-strands or the total number of strands in LptD (Fig. S3). Thus, we compared these topology predictions to independent information for evidence. Since we observed high correspondence (93 %) between ECs and the strands in the benchmark dataset, we use this approach to discriminate different topology predictions obtained for LptD and found that the topology predicted by boctopus completely overlaps with predicted ECs (except first and last strands) and has the lowest fraction (1/26) of adjacent strand-strand interactions that cannot be identified using ECs alone (Table S2). Thus 26 strands and their approximate location as predicted by boctopus and independently corroborated by EVFold-PLM were used for 3D modeling LptD.

Two runs of EC calculation are performed. In one run, EVFold_BB is employed to fold the C-terminal barrel domain (residue 200-784). In the other run, the complete LptD sequence is used to determine co-evolved residues between the barrel and the non-barrel domain. The two cysteine residues (C724 and C725) are predicted to lie on an inner loop between strands 24 and 25 (Supplementary Data). Disulphide bond formation between cysteines in the N-term domain (C31 and C173) and the C-term domain has been experimentally observed [34]. And the presence of at least one of the two disulphide bonds between the two domains is essential for the formation of LtpD/E complex with LptA [34]. The disulphide bond forming residues C173 and C725 appear within the top 20 predicted ECs between the two domains (Fig. S3 and Table S3). Three conserved proline residues (P231, P246 and P261) (Fig. 4) [35] on adjacent strands (1 – 3) are spatially close to each other (Supplementary Data). Moreover, P246 is evolutionarily coupled to 10 residues, which is the highest in terms of evolutionary couplings it appears within the top L/2 predictions (Table S4). These proline residues could induce breaks in *β*-strands allowing the lateral diffusion of substrates. Furthermore, salt-bridge forming residues D256-R277 located towards the periplasmic side also appear high (ranked 66^th^) on the ECs list (Fig. 4 and Supplementary Data). In addition, residue D256 is also evolutionarily coupled to 8 residues (ranked 4^th^ highest) within top L/2 predicted ECs (Table S4).

**Fig. 4.**
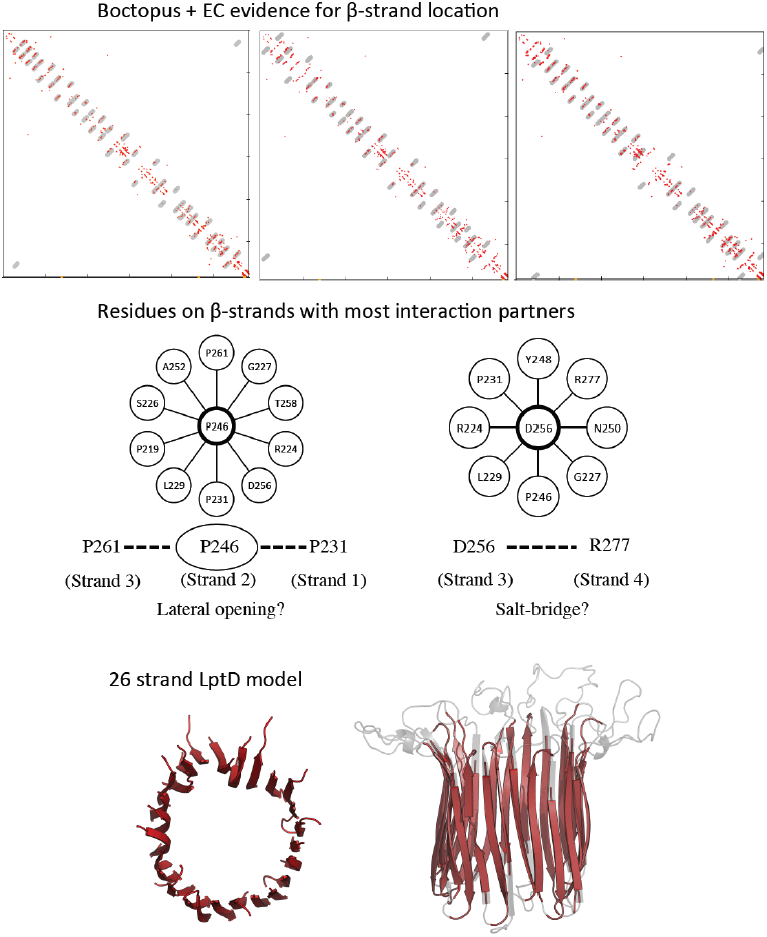
Functional insights and *de novo* predicted 3D model of LptD, a transmembrane β-barrel with no known structure. Top L/2 ECs (red) overlaid on topology predicted (gray) by boctopus, pred-tmbb and proftmb, respectively. Residue P246 located on strand 2 has the highest number of couplings (10) in top L/2 ECs and is also spatially close to P236 and P261 on adjacent strands. Residue D256 on strand 3 has 8 couplings in top L/2 predictions (ranked 4^th^). In addition, potential salt-bridge forming residues D256 are R277 are evolutionary coupled (ranked 66^th^). 26 stranded LptD model (range 200–784, strands in red) is shown in top and front view.

### Prediction of functionally important residue networks

In addition to 3D structure prediction, evolutionary couplings may identify residue networks that are functionally selected over and above those that can be identified by single column conservation alone. We calculate the evolutionary coupling count as the total number of top ranking evolutionary couplings that the residue appears in top L/2 predicted ECs. For example, in FadL protein, residue S370 with its 7 co-evolved pairs (S388, R342, A346, I345, G344, G386 and D363) is ranked high (6^th^) in terms of evolutionary couplings it appears in within top L predictions (Fig. 3 and Table S5). Site-directed mutagenesis studies have shown that S370 is required for optimal long-chain fatty acid transport in FadL [36]. ECs on the N-terminal region (A1-R42) couple with pore facing *β*-strands residues on the periplasmic side of the membrane (Fig. S4). This suggests that the N-terminal region lies in the barrel pore. For proteins with no known structure, such information will be extremely useful in understanding the physico-chemical properties of the barrel pore. Furthermore, three residues in loop 3 (P174, G176 and A185) are found to rank within top 10 on evolutionary coupling count (Fig. 3 and Table S5). Loop 3 and 4 harbor a hydrophobic groove known to be the initial low-affinity interaction site for fatty acids [12].

For FecA, interaction of plug domain residues R150 and R196 with E541 and E587 located on *β*-strands is essential not only for fixing the plug within the barrel but also plays an important role in function (Fig. 3) [37]. Residue R196 and E587 are ranked 3^rd^ and 13^th^, respectively on evolutionary coupling count list for top L predictions (Table S6). R196 appears to have co-evolved with residues D105, E587, G129, P128, A103, Q589, I149 and I127, respectively and residue E587 appears in evolutionary couplings with R150, R196, F558, E541, A611 and G556. R150 and E541 both appear in evolutionary couplings with 4 residues in top L predictions (Supplementary Data). When superimposed on the known crystal structure, the interaction network of these residues lies spatially close to each other (Fig. 3).

### Extracellular loops

In general, TMBs have long and flexible extracellular loops, some of which play a role in substrate transfer [2] and harbor sites that confer antibiotic resistance [38]. These loops are often missing in the crystal structures and rarely have many contacts with other regions in the structure as they protrude away from the membrane center. In addition, more gaps occur in loop regions as compared to the strand regions (Fig. S5). Thus, only a few high ranking ECs are predicted in the loop regions (Fig. S5). To have a better estimate of applicability of EVFold_BB in *de novo* folding the extracellular loops, we analyzed the 2969 non-trivial contacts (Methods), of which only 58 have evolutionary couplings within the top L/2 predictions (Table S7). This suggests that most of the observed non-trivial contacts might be dynamic in their nature and perhaps, have not been captured by the crystalized 3D structure.

In contrast, contacts that do appear to strongly co-evolve could indicate functionally important interactions. For example, in Q9HVS0_PSEAE (pdb: 3syb) [39], high ranked ECs are found on extracellular loops 1 to 4 (Fig. S4). Residues A35, T36 and G37 on extracellular loop 1 form EC pairs with Q95 on loop 2. In addition, ECs between residues F167, Y169, D171 and A174 on loop 3 and pore-facing residues F199, L161, R49 and E109 located on *β*-strands, suggest that loop 3 lies in the barrel pore to block the exit route from the barrel. Further, residues D212 and M213 located on extracellular loop 4 form EC pairs with K201, V202 on strand 7, respectively, and an EC pair (G224-V252) between extracellular loop 4 and 5 is also found (Table S7).

In BamA, extracellular loop 6 harbors all the 16 ECs non-trivial contacts present in top L/2 predictions (Table S7). This clearly highlights the importance of loop 6, which also contains the extensively studied VRGF/Y (660-662) motif. The residues in this motif are not only essential for the correct folding of BamA, but also for translocating proteins and cell viability [40, 41]. In the crystal structure, loop 6 interactions with strands 14-16 is mediated by R660, E698 and D721 [2]. We predict that residues on extracellular loop 6 have co-evolved with residues on *β*-strands 10 to 15 (Fig. 3 and Supplementary Data).

Interestingly, the RGF motif residues 660-662 appear high (rank 51, 30 and 66, respectively) on the list and are predicted to have co-evolved with residues on strands 12 to 15 (Supplementary Data). In addition, residue F662 in the RGF motif is ranked 4^th^ on the list of evolutionary coupling count (Table S8). Protease-sensitivity assays show that loop 6 is flexible and undergoes large conformational changes during its activation and inactivation [42]. A similar effect has also been suggested in FhaC, where the corresponding loop is captured near the periplasmic region in the crystal structure [43]. Residues G754, P719, E698, that are predicted to co-evolve with R660, are located near the periplasmic end of strand 12-14 (Fig. 3). This suggests that loop 6 undergoes a conformation change in BamA as well and residue R660 on loop 6 stabilizes this switch from the crystalized closed to open state [2].

## Discussion

We demonstrate here that evolutionary couplings together with topology predictions can be used to extract the hydrogen-bonding network between adjacent *β*-strands in TMBs. and that these constraints together with other high rank ECs from these predictions are sufficient to *de novo* fold TMBs. With enough sequence coverage, the method developed here, EVFold_BB can be used to determine the location and interaction of barrel/plug domain, which is crucial for understanding the gating mechanism of TMBs with large plug domains. For a few proteins, we show that ECs capture residues with experimentally verified functional importance. Such an approach has previously been used for helical membrane proteins and soluble proteins [23, 24], but a more rigorous analysis is needed for a reliable functional interpretation of ECs [44]. The resolution of 3D models generated by EVFold_BB is sufficient for determining the spatial location of functionally interesting regions and physico-chemical properties of the barrel domain.

All the 14 putative TMB families with an unknown structure have < 7 sequences per residue (Table S9 and Methods). With more and more genomes being sequenced every year, we anticipate that more TMB sequences will be made available soon. To automatically predict the structure of these a computational method to detect TMB domain boundaries needs to be developed to fully automate the EVFold_BB pipeline. Thereafter, a ECs based strategy could be implemented to identify transmembrane *β*-strands in putative TMBs with unknown structure. It remains to be seen if predicted ECs can be used to investigate large conformational changes that take place on multi-protein complex formation such as in FimD-CFGH complex [10]. In addition, we propose that this approach of extracting hydrogen bonds from raw predicted ECs can be generalized and extended to other *β*-sheet containing proteins as well.

## Methods

### Benchmark dataset

We started with 141 TMBs (belonging to 52 PFAM families) with 3D structures available in OPM database. Of these, 18 multi-chain TMBs were removed and 56 TMBs in 29 PFAM families were obtained after redundancy reduction at 30% sequence identity. From these, 24 PFAM families were obtained such that the alignment overlap between the two families is <= 20%. Of these 24, 18 TMBs have > 5 sequences/residue in their alignment and were chosen to benchmark EVFold_BB. These proteins cover TMBs from 8 – 24 strands. Location of the *β*-barrel domain was obtained from the PDB structure.

A list of 14 TMB families as defined in OMPD_DB were extracted from a list of 70 putative TMB families [45]. Of the 70 putative TMB families in OMP_DB, 34 have a 3D structure and 22 have a close structural homologue that can be identified using HHpred [46] (Table S9). 14 putative TMB families have no known 3D structure for a representative sequence. All these families have < 7 sequences / residues in their alignment, which is at the low end of number of sequences / residues that EVFold_BB has been benchmarked on. From this list, LptD is chosen as an interesting protein as substantial experimental information is available in the literature [29, 34, 47, 48].

### Prediction of Evolutionary couplings from multiple sequence alignments

Multiple sequence alignments for all proteins were generated using jackhmmer (version – 3.1) [49] against the UniProt database. For all proteins, three iterations were performed at an E-value of 10^−2^ to ensure maximum number of sequences. For full length LptD (residues 15-784), an E-value of 10^−10^ was chosen to optimize the number of sequences but also have sufficient sequence coverage. A global statistical inference method based on pseudo-likelihood maximization [27] as implemented in EVFold (EVfold.org) [24] is employed to extract direct interactions from all the observed correlations in a MSA. A ranked list of ECs is obtained by taking the norm of the matrix of couplings and adjusting for phylogenetic bias using average-product correction [27].

### Topology prediction using boctopus2.0

A non-redundant dataset (< 30% sequence identity) of 36 TMBs with known structures along with transmembrane *β*-strand boundaries was curated from the OPM database [30] (Table S10). All residues in the dataset were labeled as – outer loop (o), inner loop (i), *β*-strand pore facing (P), *β*-strand lipid facing (L). The position specific scoring matrix (PSSM) obtained using three iterations of hhblits (version - 2.0.13) [50] against nr database (nr20_12Aug11) is used as the input to four separate support vector machines (SVMs) that were trained to predict the per-residue location. Together with secondary structure prediction using PSIPRED [51], a per-residue profile is generated and used as input to a hidden Markov model to predict the overall topology. Boctopus2.0 is trained based on a 10-fold cross-validation where all proteins belonging to the same family were put together. For the barrel domain, boctopus2.0 predicts the correct topology for 32 out of 36 proteins in the benchmark dataset. Number of strands is correctly predicted for 34/36 proteins except 1i78 and 2qdz (Table S10). For PORP_PEASE (pdb: 2o4v), two extra strands are predicted outside the barrel domain. Topologies for 5 proteins not in the boctopus2.0 dataset (3syb, 4k3c, 4e1t, 3ohn, 2jk4) are predicted using trained boctopus2.0 (Table S10). For INVA_YERPS (pdb: 4e1t) and VDAC1_HUMAN (pdb: 2jk4), 2 and 1 extra strand are predicted outside the barrel boundary, respectively. For Q93PM2_HAEDC (pdb: 4k3c), FECA_ECOLI (pdb: 1kmp) and ESTA_PEASE (pdb: 3kvn), the non-barrel/barrel boundary is predicted by using “P” and “L” probabilities averaged over a window size of 50 residues (Fig. S6). Region with the average value > 0.6 was classified as barrel.

### Determination of ***β***-strand registration

The residues in predicted ECs are annotated with their strand location and face status (pore-facing or lipid-facing) obtained from boctopus2.0. A list of ECs, where the residue pairs lie on adjacent strands, is generated and top L (where L = length of the protein) ECs are extracted. Predicted *β*-strands are taken in a pairwise manner and shifted +/-3 residues with respect to each other to generate alternate pairs. For each configuration, EC strength of all pairs is summed. Residue pairs that do not face in the same direction are penalized by –1. The shift with the highest EC strength is chosen for that strand pair. Hydrogen bonds are put on alternate residues such that the dyad repeat pattern and the right-handed twist of TMBs is maintained throughout the barrel [52]. This is done by placing hydrogen bonds only on porefacing residues if the paired strands traverse from up (periplasmic to extracellular) to down (extracellular to periplasmic) and on lipid-facing residues if the paired strands traverse from down to up. For comparison, observed hydrogen bonds are extracted from known 3D structure, residues on adjacent strands are considered hydrogen bonded if the distance between their N-O atoms is < 3.4 Å. The cutoff is based on distribution of all adjacent N-O bond distances such that most hydrogen bonds are included.

### Resolution of unlikely constraints

The *β*-strand boundaries predicted by boctopus2.0 are superimposed on the secondary structure predicted by PSIPRED [51]. Location of predicted *β*-strands and loops is used to filter out constraints that are considered unviable as described by Marks *et al.* [24]. In addition, ECs are annotated to be “strand-strand” or “non strand-strand” based on if both the residues are located within predicted *β*-strand boundaries or not.

### Distance constraints from predicted hydrogen bonds and ECs

Default values for distance constraints are derived from the distance distribution observed in transmembrane *β*-strands with a known 3D structure. Distance constraints are put on side-chain heavy atoms, O-N, N-O and CA-CA atoms for residues that are predicted to be hydrogen bonded. For other ECs between non strand-strand residues, distance constraints are put on side-chain heavy atoms only. Secondary structure distance constraints are put on O-O, N-N, CA-CA, CB-CB, O-N, CA-O atoms and dihedral angles are constraint with default values for an anti-parallel *β*-sheet (Table S11 and S12).

### *de novo* folding using CNS

Only a small number of models (M = 50) per set of constraints used are generated. We start with applying only predicted hydrogen-bond constraints on adjacent strands to fold TMBs. Other EC constraints are included in steps (S =10) up to L/2, where L is the length of protein. In addition, constraints are put to maintain the predicted secondary structure. Distance constraints are used in CNS [31] to *de novo* fold TMBs. CNS uses a distance geometry protocol followed by simulated annealing to satisfy the input constraints. All folding predictions start with a fully extended polypeptide chain. A square potential well implemented in CNS is used to penalize constraint violations. After annealing, a short two-stage energy minimization step is employed to relax generated structures and add hydrogen bonds. For example, for Q9RP17_NEIME (pdb: 1p4t) that has a length of L =160, (1 + L/2*S)*M = 450 models are generated. To facilitate folding of ∼120 residues FECA_ECOLI non-barrel plug domain, models are generated starting with at least 60 constraints involving the plug domain and up to L/2 non strand-strand constraints, where L is the length of the entire protein with both domains.

### Blinded model ranking

Generated models are ranked based on the score obtained by summing the fraction of hydrogen bond constraints satisfied in the barrel region of the generated model and β-twist score as defined by Marks *et al.* [24]. A hydrogen bond constraint is considered satisfied if the distance between the N-O atoms is 2.9 +/-0.3 Å. The twist score and fraction of hydrogen bond constraints satisfied are normalized before addition. FECA_ECOLI (barrel+plug) models are ranked based on the twist score and fraction of hydrogen bond constraints satisfied in the barrel domain and the number of constraints satisfied in the plug domain and plug/barrel interface. In addition, the barrel region is compared to the corresponding region in the known structure using TM-score [53] to access the model quality when a crystal structure is available.

### Estimation of Non-trivial contacts

Non-trivial contacts are defined as residue pairs that are at least 4 residues away from their nearest full *β*-strand (membrane boundary + DSSP [54]) when present on adjacent strands. When there is at least one transmembrane *β*-strand between the residue pair, then the minimum distance of +2 residues from the full *β*-strand is considered. Furthermore, self-loop contacts are excluded.

### Evolutionary coupling count

For each residue, we defined evolutionary coupling count as the sum of EC scores of evolutionary couplings a residue occurs in within top L predictions. The general idea is that a functionally important residue has stronger couplings resulting in high scoring ECs with more interaction partners. Thus a high evolutionary coupling score could signify a functionally important site.

## Acknowledgement

DSM, CS and SH are supported by NIH award R01 GM106303. AE is supported by grants from the Swedish Research Council (VR-NT 2012-5046, VR-M 2010-3555), SSF (the Foundation for Strategic Research) and Vinnova through the Vinnova-JSP Program.

## NOTE

During writing of this manuscript two papers describing the 3D crystal structure of LptD with 26 *β*-strands have been published*. No information from those publications was used in this study and the PDB co-ordinates (4Q35) and (4N4R) were not available at the time of submission of this manuscript. *Dong *et. al*, “Structural basis for outer membrane lipopolysaccharide insertion”, Nature 2014 and Qiao *et al*., “Structural basis for lipopolysaccharide insertion in the bacterial outer membrane”, Nature 2014.

## List of Supplementary Figure Legends

**Fig. S1** – Predicted contact maps using EVFold-PLM. Predicted ECs (red) on observed crystal contacts (<= 5Å) (gray), gaps in 3D structure in pdb file (blue).

**Fig. S2** – Location of gaps in MSA. Predicted (pink) and observed (green) β-strand boundaries overlap. Percentage of gaps (-) per column in the multiple sequence alignment is higher in loop regions. In the case of Q9HVD1_PSEAE and PORP_PSEAE, columns corresponding to the first and the last strands have > 50% gaps in the MSA.

**Fig. S3** – Predicted LptD topologies using three different prediction methods and contactmap showing inter-domain interactions. Disuphide bond C173-C725 is highlighted.

**Fig. S4** –

Fig. S4A – ECs suggest interaction between N-term helix and barrel interior in FADL_ECOLI (pdb: 1t16). Residues on the N-term helix are evolutionary coupled with residues in the barrel pore region (red).

Fig. S4B – ECs in loops reveal functional sites. Q9HVS0_PSEAE (pdb: 3syb) has 11 non-trivial contacts in long loops that make non-trivial contacts with residues on β-strands and block the pore exit.

**Fig. S5** –

Fig. 5A – Distribution of columns with more than 50% gaps show that loops (blue) have more and larger gaps than strands (pink).

Fig. 5B – Only a few ECs in loop regions in top L. Predicted ECs are superimposed on the TMBs structures aligned to membrane (OPM_DB). Distribution of ECs is higher around the membrane center that contains the β-strand that form the barrel pore and starts decreasing +/13 Å on either side (loop region). There are fewer ECs in loop-loop region (black) as compared to other regions.

**Fig. S6** – Boctopus domain boundary detection

